# Recent transposable element bursts triggered by insertions near genes in a fungal pathogen

**DOI:** 10.1101/2022.07.13.499862

**Authors:** Ursula Oggenfuss, Daniel Croll

**Affiliations:** Laboratory of Evolutionary Genetics, Institute of Biology, University of Neuchâtel, 2000 Neuchâtel, Switzerland

## Abstract

The activity of transposable elements (TEs) contributes significantly to genome evolution. TEs often destabilize genome integrity but may also confer adaptive variation in phenotypic traits. De-repression of epigenetically silenced TEs often initiates bursts of transposition activity that may be counteracted by purifying selection and genome defenses. However, how these forces interact to determine the expansion routes of TEs within a species remains largely unknown. Here, we analyzed a set of 19 telomere-to-telomere genomes of the fungal wheat pathogen *Zymoseptoria tritici*. Phylogenetic reconstruction and ancestral state estimates of individual TE families revealed that TEs have undergone distinct activation and repression periods resulting in highly uneven copy numbers between genomes of the same species. Most TEs are clustered in gene poor niches, indicating strong purifying selection against insertions near coding sequences. TE families with high copy numbers have low sequence divergence and strong signatures of defense mechanisms (*i*.*e*., RIP). In contrast, small non-autonomous TEs (*i*.*e*., MITEs) are less impacted by defense mechanisms and are often located in close proximity to genes. Individual TE families have experienced multiple distinct burst events that generated many nearly identical copies. We found that a *Copia* element burst was initiated from recent copies inserted substantially closer to genes compared to older insertions. Overall, TE bursts tended to initiate from copies in GC-rich niches that escaped inactivation by genomic defenses. Our work shows how specific genomic environments features provide triggers for TE proliferation.

## INTRODUCTION

Transposable elements create novel insertions by duplication or relocation of existing copies in the genome. Left unchecked, TEs may proliferate in the genome (*i*.*e*., burst), leading to genome expansions, increased ectopic recombination and the potential of deleterious insertions into coding and regulatory regions (Petrov *et al*., 2003). Defense mechanisms have evolved to reversibly inactivate (*i*.*e*., silence) or irreversibly mutate TEs (Daboussi & Capy, 2003; Lisch & Bennetzen, 2011). Defenses against TEs include histone modifications, cytosine and chromatin methylation, small RNA based silencing or KRAB zinc finger based transcriptional silencing (Lisch, 2009; Jacobs *et al*., 2014; Yang *et al*., 2017; Schmitz *et al*., 2019). Some TEs have the ability to regulate their own expression through small RNAs (Rebollo *et al*., 2012). In certain ascomycete fungi, TEs are also targeted by repeat-induced point mutations (RIP) during sexual recombination (Galagan & Selker, 2004; Gladyshev & Kleckner, 2017; van Wyk *et al*., 2021). RIP introduces CpA to TpA mutations in all copies of a duplicated sequence. RIP generally decreases GC content and typically leads to loss-of-function at targeted loci. In the ascomycete *Neurospora crassa*, a few generations of sexual recombination are sufficient to degenerate TE copies (Wang *et al*., 2020b). TE expansions are also counterbalanced by deletion via ectopic recombination, purifying selection and genetic drift (Charlesworth & Charlesworth, 1983; Devos *et al*., 2002; Petrov *et al*., 2003). Defense mechanisms against TEs may be weakened under stress conditions or lost over evolutionary time scales (González *et al*., 2008; Horváth *et al*., 2017; Lorrain *et al*., 2020). In the absence of bursts, most TE families are expected to reach a plateau in copy numbers determined by a balance between insertion and deletion rates (Charlesworth & Charlesworth, 1983; Petrov *et al*., 2003; Le Rouzic & Capy, 2005). A loss of control over TEs can lead to de-repression and rapid copy number amplifications, which may have dramatic impacts on the integrity of the genome, and is often associated with speciation events (Belyayev, 2014). How and why specific TE copies initiate bursts remain unclear; however, an increase in copy number of previously de-repressed TEs or an introduction of new TE families into a species often occur in tandem with genomic defenses being activated (Le Rouzic & Capy, 2005).

How TE insertion reshape the genomic landscape depends on the fitness effects of new insertions (Sigman & Slotkin, 2016). Generally, strong purifying selection acts against most new insertions in plant, animal and fungal genomes (Cridland *et al*., 2013; Stuart *et al*., 2016; Lai *et al*., 2017; Stritt *et al*., 2017; Oggenfuss *et al*., 2021). However, some niches in the genome tolerate or even favor new TE insertions (Kremer *et al*., 2020; Stitzer *et al*., 2021). Some TE superfamilies have preferred insertion sites characterized by specific motifs or other sequence characteristics (Bridier-Nahmias *et al*., 2015; Sultana *et al*., 2017). Insertion site preferences may lead to genomic compartmentalization resulting in the generation of large gene-sparse niches of nested TEs. Many fungal plant pathogens accumulate TEs in gene-sparse niches and accessory chromosomes, and are often co-located with pathogenicity-associated genes (Rouxel *et al*., 2011; Raffaele & Kamoun, 2012; Van Dam *et al*., 2017; de Freitas Pereira *et al*., 2018; Torres *et al*., 2020). Additionally, few TE insertions can provide a positive impact and will be under positive selection (González *et al*., 2008; van’t Hof *et al*., 2016; Horváth *et al*., 2017; Zhang *et al*., 2019). As a consequence of insertion site bias, natural selection and defense mechanisms, most TE families are unevenly distributed across the genome (Bourque *et al*., 2018). To what degree TEs have adapted to specific niches of the genome or simply reflect the outcome of purifying selection remains largely unclear.

Low divergence among TE sequences within a given family generally indicates that a burst of activity has occurred recently (Lerat *et al*., 2003). Hence, phylogenetic analyses of TE copies can reconstruct the evolutionary history of the TE family. Analogous to viral birth-death models, bursts of transpositions should leave distinct marks of short internal branches in phylogenetic trees (Volz *et al*., 2013; Blumenstiel, 2019). Transposition bursts are characterized by a most recent common ancestor likely reflecting the copy initiating the expansion. In contrast, copies with long terminal branches have likely been silenced. Such TEs will initially remain in populations as remnants, accumulate mutations and ultimately degenerate. Reconstructing the sequence of events leading to transposition bursts is often challenged by the difficulty in recovering all copies of a TE within a given species. The difficulty stems from the incomplete nature of many genome assemblies and the fact that insertions are not fixed in populations. Recent bursts of TEs were found in large collections of rice genome resequencing datasets showing that highly active elements can be successfully identified (Lu *et al*., 2017). Yet recovering full-length copies of TEs that resulted from transposition bursts remains challenging without high-quality genome assemblies.

Many filamentous fungal plant pathogens have compact genomes and a high degree of genomic compartmentalization into TE dense and gene dense niches, thus making them amenable to TE analyses (Frantzeskakis *et al*., 2019; Torres *et al*., 2020). Genes present in TE rich niches are often dispensable for survival but may locally play an important role in virulence on the host (*i*.*e*., effectors) (Raffaele & Kamoun, 2012; Croll & McDonald, 2012; Faino *et al*., 2016; Torres *et al*., 2020). *Zymoseptoria tritici* is an important fungal plant pathogen on wheat that co-evolved with its host (Stukenbrock *et al*., 2007). TEs cover 16.5-24% of the genome and TE bursts are associated with not only incipient genome size expansions (Oggenfuss *et al*., 2021) but with the emergence of adaptive traits. For example, insertions of multiple TEs in the promoter region of a major facilitator superfamily transporter led to multidrug resistance in *Z. tritici* (Omrane *et al*., 2015, 2017; Mäe *et al*., 2020). Furthermore, increased activity of a DNA transposon reduces asexual spore production, melanization and virulence (Krishnan *et al*., 2018; Meile *et al*., 2018; Wang *et al*., 2021). Some populations of the pathogen have likely lost *dim2*, an essential gene of the RIP machinery (Möller *et al*., 2020; Lorrain *et al*., 2021). Cytosine methylation was also lost after a duplication event of the *MgDNMT* gene and subsequent RIP mutations rendered all copies non-functional (Dhillon *et al*., 2010, 2014). Histone modifications and small RNAs likely contribute to silencing of TEs (Schotanus *et al*., 2015; Kettles *et al*., 2019; Habig *et al*., 2021). The TE repertoire includes 304 TE families based on an analysis of 19 completely assembled genomes (Badet *et al*., 2020). A subset of TEs show evidence for de-repression during plant infection likely connected to bursts of TE proliferation (Fouché *et al*., 2020; Badet *et al*., 2020; Oggenfuss *et al*., 2021).

Here, we retraced the invasion dynamics of young TEs in the recent history of *Z. tritici* to determine triggers of TE expansions. We established near total evidence for all copies of multiple TE families by combining information from 19 telomere-to-telomere genome assemblies. We use phylogenetic reconstruction of ancestral TE states to identify genomic niches of TE activation and proliferation. The comprehensive view of TE invasion routes establishes that insertions near genes act as triggers for copy number expansions. Escape of genomic defenses including silencing is a likely the major driver of TE dynamics.

## RESULTS

### TRANSPOSABLE ELEMENT DIVERSITY WITHIN *Z. TRITICI*

We analyzed 19 chromosome-level genomes to comprehensively map the genome-wide distribution of TE families in the fungal pathogen *Z. tritici* (Figure 1A; (Badet *et al*., 2020)). Genome size and TE content vary considerably among individuals and show a positive correlation. The genomes of an Australian and Iranian isolate have the highest TE content (24%) and lowest TE content (18.1%), as well as the largest genome (41.76 Mb) and smallest genome (37.13 Mb), respectively (Figure 1B, Supplementary Table S1). We focused on TE families with at least 20 copies, which collectively have a total of 24,520 copies across all analyzed genomes. Half of the copies belong to DNA transposons (*n* = 104 families) and half to retrotransposons (*n* = 59 families; Supplementary Table S2) without meaningful differences among genomes (Figure 1C; Supplementary Figure S1). Most TE insertions are singletons or are at low frequency (Figure 1D). Only few TE insertions are fixed among genomes (*n* = 122; Figure 1D) and consist predominantly of MITEs.

**Figure 1:**
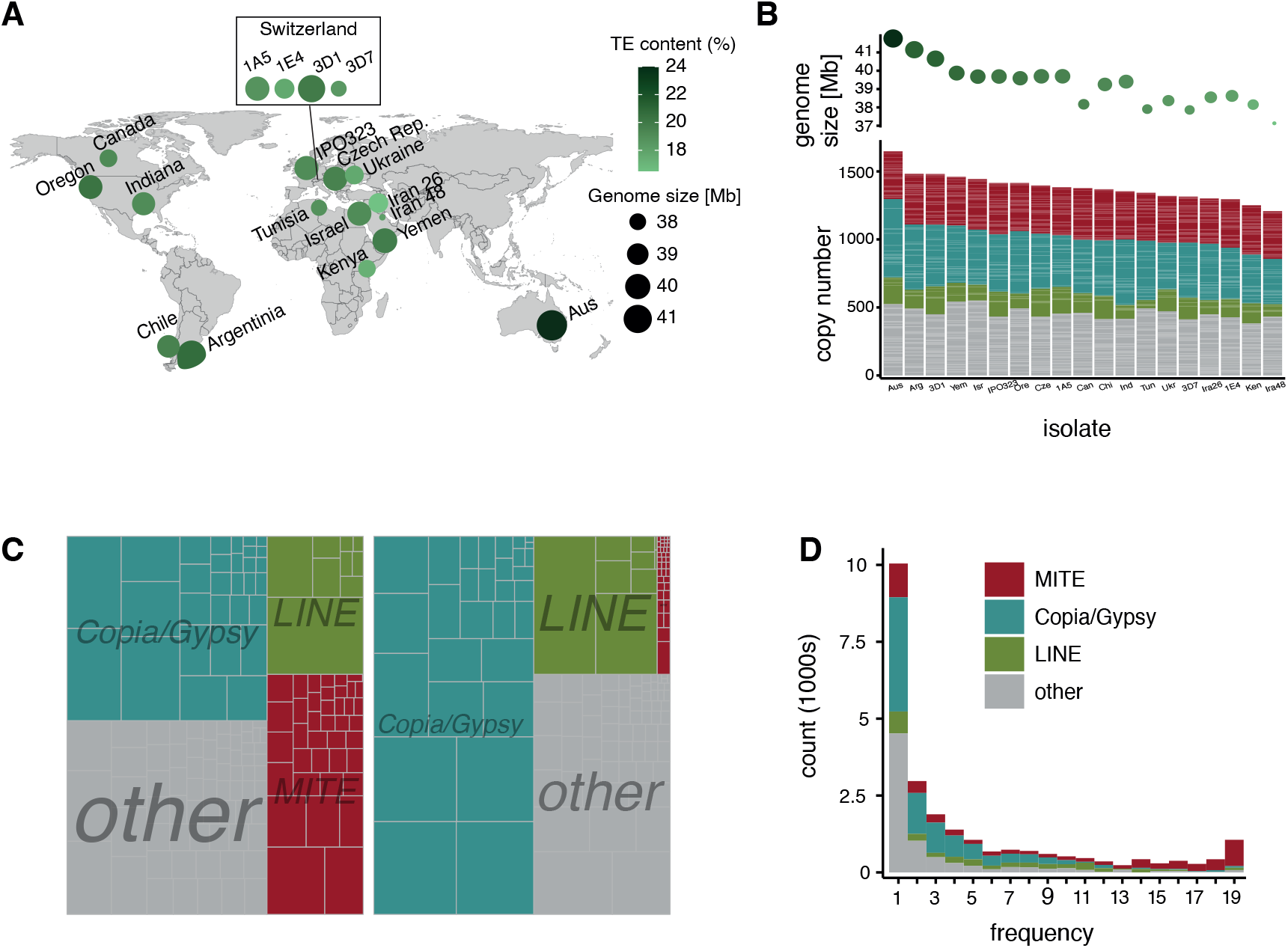
Transposable element (TE) distribution in 19 telomere-to-telomere genomes of *Zymoseptoria tritici*: (A) Origin of isolates used for genome analyses. Circle size indicates the genome size, the green shade indicates the TE content. (B) Genome size and TE copy number per isolate. The colors indicate MITEs (miniature inverted repeat transposable elements, small non-autonomous DNA transposons corresponding to several TE superfamilies), RLC and RLG (two superfamilies belonging to LTR) and LINE. (C) Copy numbers of TEs (left) and total length (right) in all 19 genomes. Smaller boxes correspond to TE families. (D) Allele frequencies of TEs at orthologous insertion loci among genomes. TEs were defined as orthologous if they were located between the same set of orthologous genes.

### TRANSPOSABLE ELEMENT INSERTION NICHES HAVE LOW GENE CONTENT

We analyzed 5kb niches centered around TE insertions (Figure 2A). For the reference genome IPO323, we found no accumulation of TE insertions in niches with marks of open chromatin (*i*.*e*., euchromatin). A small subset of TE insertions overlaps with obligate heterochromatin marks (Figure 2A). Across all 19 genomes, we found that most TEs are located on core chromosomes (*i*.*e*., chromosomes shared among all isolates), but TE insertions are at higher density on accessory chromosomes (Figure 2B). The majority of TE copies are located in niches with a GC content below 50%, with the exception of MITEs (average of 72.0% GC; Figure 2B). Only a small subset of TE copies is in niches overlapping a gene or a subtelomeric region (Figure 2B). We found no overall association between TE insertions and large RIP-affected regions. However, most RLC, RLG and LINEs are located inside and most MITEs outside of large RIP affected regions (Figure 2B).

**Figure 2:**
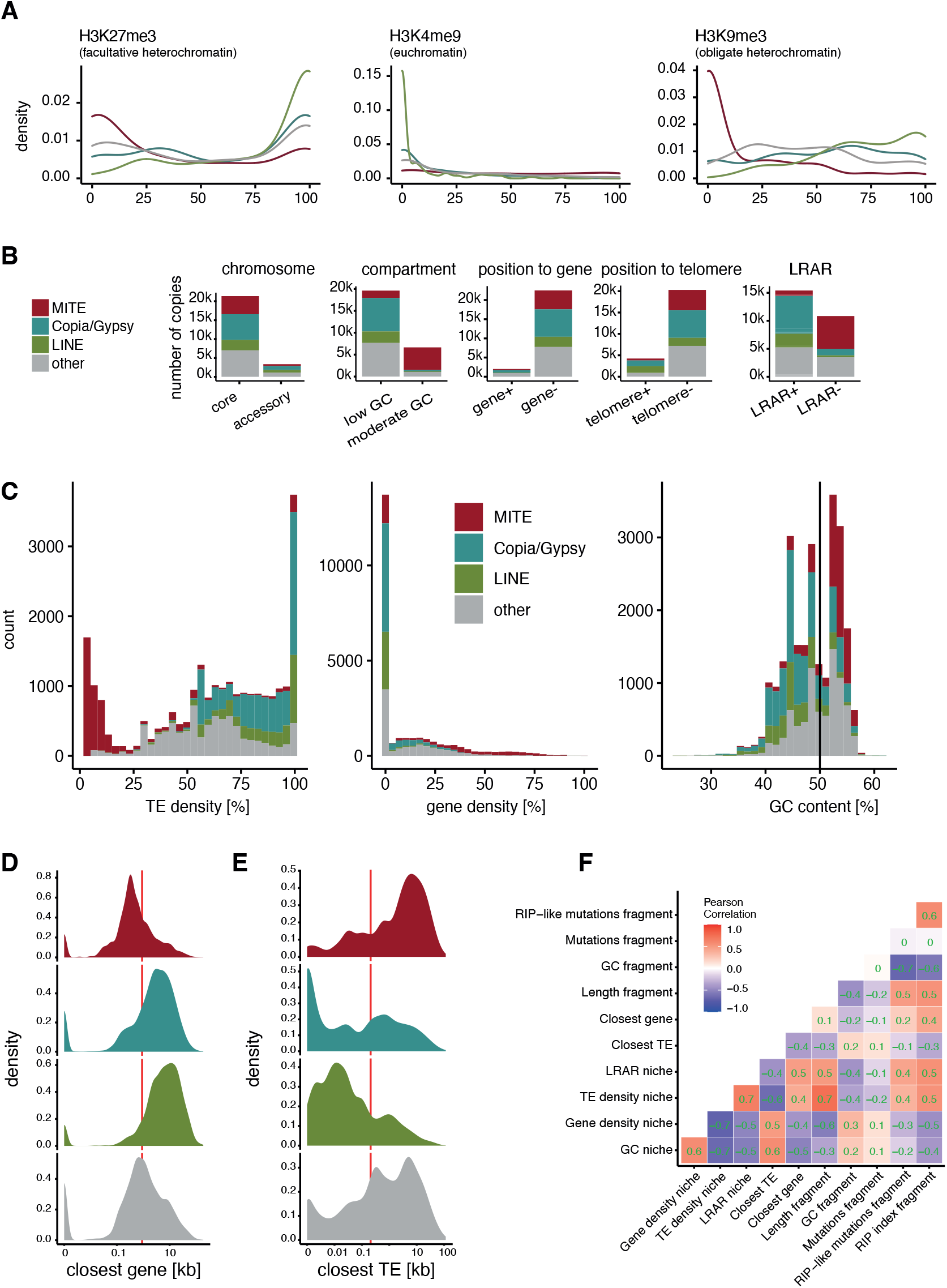
Characteristics of TE insertion niches across the genome: (A) Proportional overlap of H3K27me2, H3K4me9 and H3K9me3 histone methylation marks with TE insertion niches in the reference genome IPO323. Colors indicate the group of TE. (B) TE insertion sites between core and accessory chromosomes, TE insertions into niches with a moderate (≥ 50%) or low (< 50%) GC content, TE insertions into regions annotated as genes, TE insertions into subtelomeric region and TE insertions into large RIP affected regions. (C) Overlap of TE content, gene content and GC content with TE insertion niches. (D) Distribution of the distance to the closest gene in MITEs, RLC/RLG, LINE and other TEs. The red line indicates the mean distance. (E) Distances to the next TE MITEs, RLC/RLG, LINE and other TEs. The red line indicates the mean distance. (F) Spearman correlation matrix of 11 characteristics of TE insertion niches and TE copy characteristics. Dark red indicates strong positive correlation, dark blue indicates strong negative correlation of two characteristics.

More than one third of TEs are inserted into niches with more than 80% TE content. In contrast, MITEs are preferentially inserted into TE poor niches (Figure 2C). GC content in TE insertion niches varies between 25-60%. We found more than one third of TE insertions 1-10kb away from the next gene, with MITEs being on average closer (Figure 2D). TE insertions are often close to RLC, RLG and LINEs copies (902 and 2431 bp average distance, respectively). MITEs generally are at a distance 8,037 bp from the next TE on average (Figure 2E). Overall, TE density is negatively correlated with gene density and GC content (Figure 2F). Longer TE copies tend to be located in already TE rich niches.

### RECENT ACTIVITY OF HIGH-COPY TRANSPOSABLE ELEMENT FAMILIES

Recently active TE families typically carry a high number of similar TE copies in the genome. We first filtered for a subset of TE families with more than 100 copies in all 19 genomes combined. The 61 retained TE families predominantly include MITEs (*n* = 12) as well as RLC and RLG (*n* = 5 and 11, respectively; Supplementary Figure S2A). We find that high-copy TE families tend to also have more variable copy numbers among 19 genomes, indicating ongoing activity of individual families (Figure 3A). The GC content of high copy TE families generally approaches the genome-wide GC content, with the exception of MITEs that have an overall higher GC content around 52%, and an RLC_Deimos family where most copies have a GC content below 40% (Figure 3B). Full-length TE copies range from 218-13,907 bp, with the shorter copies belonging to the non-autonomous MITEs lacking coding regions (Figure 3C).

**Figure 3:**
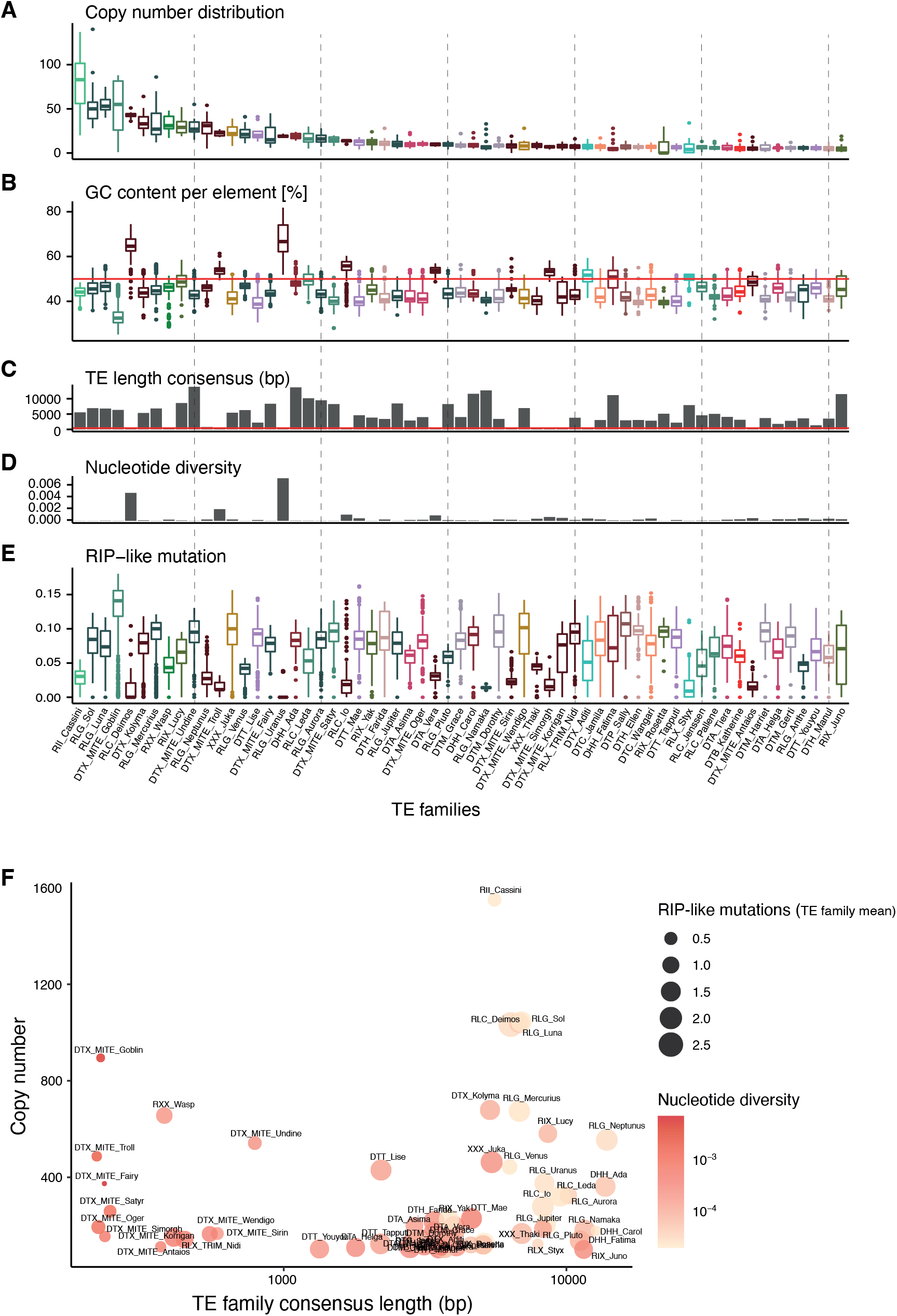
Characteristics of high-copy TE families: TE families are ordered from the highest copy numbers to lowest copy numbers (right) in all 19 analyzed genomes combined. (A) Total copy numbers. (B) Copy numbers per TE family. (C) GC content distribution per TE family. (D) Length of the consensus sequence corresponding to the full-length consensus sequence excluding nested TEs or partial deletions. (E) Nucleotide diversity of the TE family. (F) Number of RIP-like mutation (CpA<->TpA/TpG<->TpA) per TE copy, corrected for the length of the TE. (G) Long (> 0.00001; red) and short (≤0.00001; blue) terminal branch lengths of individual copies characterizing two classes of divergence times. (H) Correlation between copy numbers and consensus sequence lengths for TE families. Circle size corresponds to the mean number of RIP-like mutations and the color indicates the nucleotide diversity.

To estimate the relative age of episodes with TE copy number increases, we calculated the nucleotide diversity for each TE family. The highest copy number TEs have very low nucleotide diversity consistent with recent proliferation in the genome. MITEs tend to have higher nucleotide diversity at similar copy numbers compared to other TE families (Figure 3D, F). MITEs are also less affected by RIP (Figure 3E, F). Terminal branch lengths of individual TE copies are a further indication of the age of recent transposition. Copies of MITEs tend to have overall short terminal branch lengths compared to other TEs (Supplementary Figure S2B). The short length of MITEs might constrain the potential to accumulate mutations compared to longer TEs. Consistent with this, many MITEs show long internal branch lengths between distinct clades characterizing independent bursts (Supplementary Figure S3). Overall, TE families with high copy numbers and long consensus sequences show lower nucleotide diversity, however, most of the mutations are RIP-like mutations (Figure 3H). Our findings show that recent bursts are characterized by high copy numbers, but genomic defenses and the length of TEs create complex outcomes for individual TE families.

### TRANSPOSABLE ELEMENT EXPANSION ROUTES IDENTIFY GENOMIC NICHES OF PROLIFERATION

To disentangle factors influencing recent TE bursts, we first calculated genetic distances among copies (Figure 4A). Most TE families show their highest activity in a similar, recent age range. We found two TE families with ongoing activity (Styx and Thrym). Among the high copy TE families, the RII_Cassini has been most recently active. RLG_Luna, RLG_Sol, RIX_Lucy and RLC_Deimos have undergone earlier bursts with both RLC_Deimos and RLG_Luna showing indications of multiple episodes of rapid proliferation. To reconstruct TE expansion routes in the genome, we built phylogenetic trees, rooted using TE copies found in *Zymoseptoria* sister species. Using tree reconstruction, we assessed for each TE copy whether characteristics of the TE sequence and the genomic niche evolved from the parental node in the tree. We used ancestral state reconstruction to identify niche and sequence features of all internal nodes on each family’s tree. We found increases from inferred ancestral states to GC content of the TE, as well as the GC and gene content of the genomic niche (Figure 4B). While most TEs are located in a different chromosome compared to the parental node, new insertions typically remain on core chromosomes (65.4%) or switched from an accessory to a core chromosome (21.5%). We found that more than half of the insertions remain in isochores of low GC content (58.4%) or jumped from moderate to low GC content (20.5%). Additionally, more than half of all TE insertions either remain in large RIP-affected regions or jumped into such regions (20.7%).

**Figure 4:**
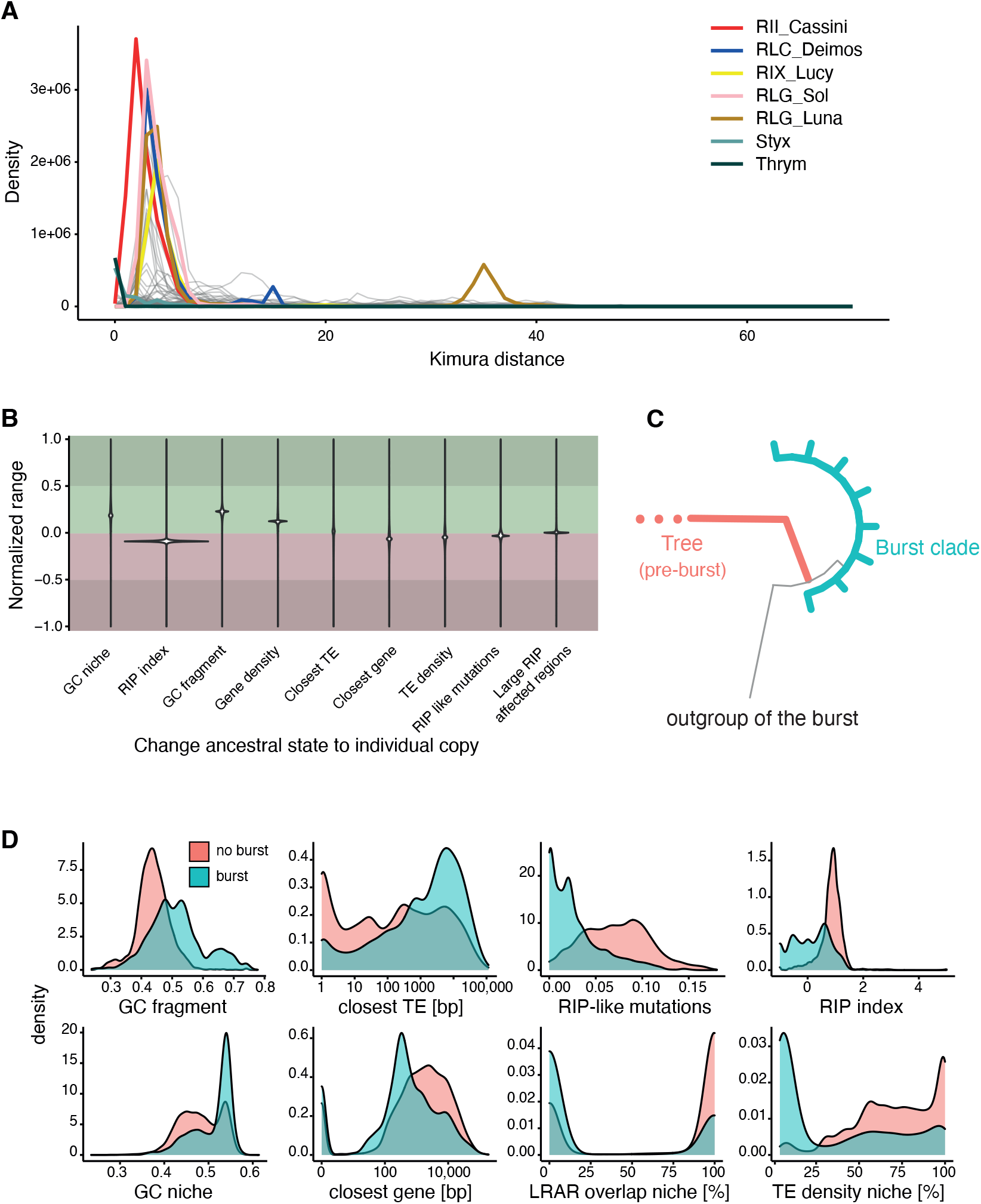
Genomic origin and features of TE transposition bursts: (A) Repeat landscape of the TE families with the highest copy numbers. Colors indicate the highest copy numbers (RII_Cassini, RLC_Deimos, RLG_Sol, RLG_Luna), TEs with multiple bursts (RLC_Deimos, RIX_Lucy and RLG_Luna) or very recent burst (Styx, Thrym). (B) Normalized range of characteristics of TE copies and their genomic niche compared between ancestral states and derived copies. Green indicates an increase and red a decrease in individual features. (C) Scheme of the definition of burst and burst outgroups based on phylogenetic trees of TE families. Green indicates the copies of a burst with low terminal branch lengths and the red outgroup indicates the closest related sister branch of a burst. (D) Distribution of niche and TE copy characteristics of copies belonging to a burst clade (red) compared to all other copies (blue).

We identified individual bursts within TE families by retrieving clades of highly similar sequences distinct from other sequences in the tree (Figure 4C). We defined the outgroup of individual bursts as the most likely parental copy that preceded the burst. Overall, around 50% of TE families experienced at least one recent burst and 10% (*n* = 32) of all TE families revealed several bursts. Copies resulting from individual bursts are often found only in a subset of the analyzed genomes of the species, consistent with TE bursts being recent. In MITE families, a large proportion of all copies likely originate from a recent burst. To identify general properties of TE bursts, we compared copies included in a burst with all other copies of from that TE family (Figure 4D). Burst copies generally have a higher GC content, less RIP-like mutations, are closer to genes and are more distant to other TEs compared to non-burst copies (Figure 4D). We found also that burst copies tend to occupy genomic niches with lower TE density, overlap less with large RIP-affected regions and have a higher GC content.

### NICHE CHARACTERISTICS OF TRANSPOSABLE ELEMENT ACTIVATION

We focused on five TE families with particularly high copy numbers and evidence for large bursts to identify drivers of expansions (*i*.*e*., LINE/*I* RII_Cassini, LTRs RLG_Luna RLG_Sol, and RLC_Deimos, and the *MITE* DTX_MITE_Goblin). The recent burst of RII_Cassini has created copies with a higher GC content carrying less RIP-like mutations (Figure 5). RII_Cassini has likely undergone six individual bursts of which two are very recent. Copies of these two bursts are each exclusively found in a single genome (isolates from Australia and Canada, respectively). Recent burst copies of RII_Cassini have higher GC content and lower numbers of RIP-like mutations compared to all other copies (Figure 6). However, most copies are inserted into niches that presently show strong signatures of RIP, moderate GC content and distant from coding sequences. Similarly, during the recent burst of RLC_Deimos, new copies were mostly inserted outside of large RIP affected regions and closer to genes. Burst copies themselves are also of higher GC content and show only few RIP-like mutations (Figure 5). RLC_Deimos has undergone a single large burst with copies carrying nearly no RIP-like mutations contrasting with most older copies being heavily affected by RIP-like mutations (Figure 7). Burst copies show an increase in GC content and are located in niches devoid of RIP-like mutations compared to all other copies. Interestingly, one of the putative copies leading to the burst of RLC_Deimos is found only in a single genome and inserted into a gene encoding an alpha/beta hydrolase. In contrast, RLG_Sol and RLG_Luna show no evidence of recent activity (Figure 5). DTX_MITE_Goblin general has high copy numbers with similar numbers and in orthologous positions among genomes, consistent with TE activity predating speciation. The expansion of DTX_MITE_Goblin is characterized by a high number of bursts (Supplementary Figure S3). Copies from individual bursts are typically shared among all genomes. Despite high nucleotide diversity, few mutations were generated by RIP. The DTX_MITE_Goblin family might consist of older insertions shared between genomes, that are not affected by RIP. Taken together, high copy number TEs tend to have very recent burst origins, a high GC content and an ability to evade genomic defenses.

**Figure 5:**
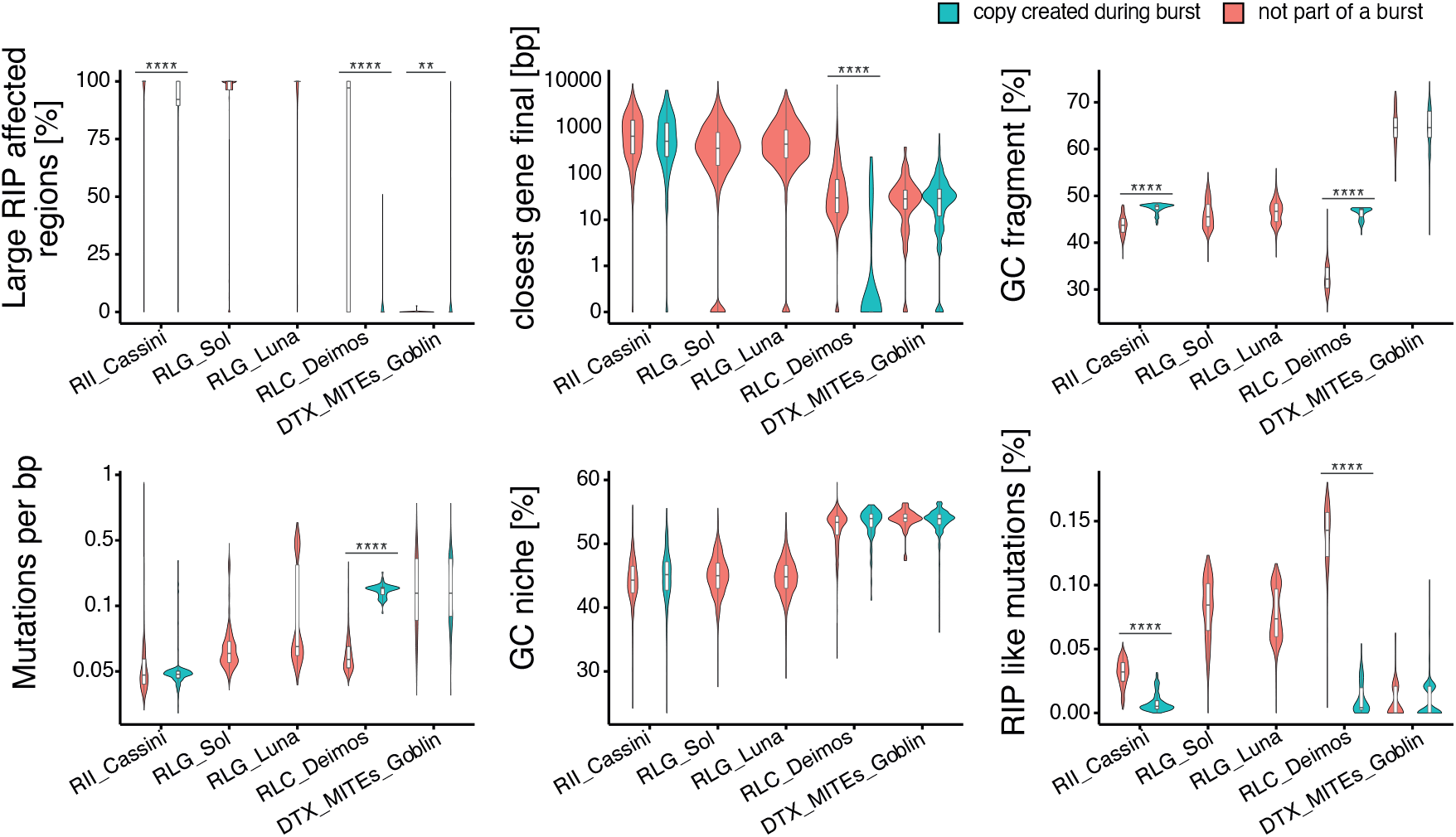
High copy number TE families and characteristics of burst initiation: Comparison of copies in bursts (red) and all other copies (blue) for the five TE families with the highest copy numbers. The TE families RLG_Luna and RLG_Sol have no copies assigned to burst clades.

**Figure 6:**
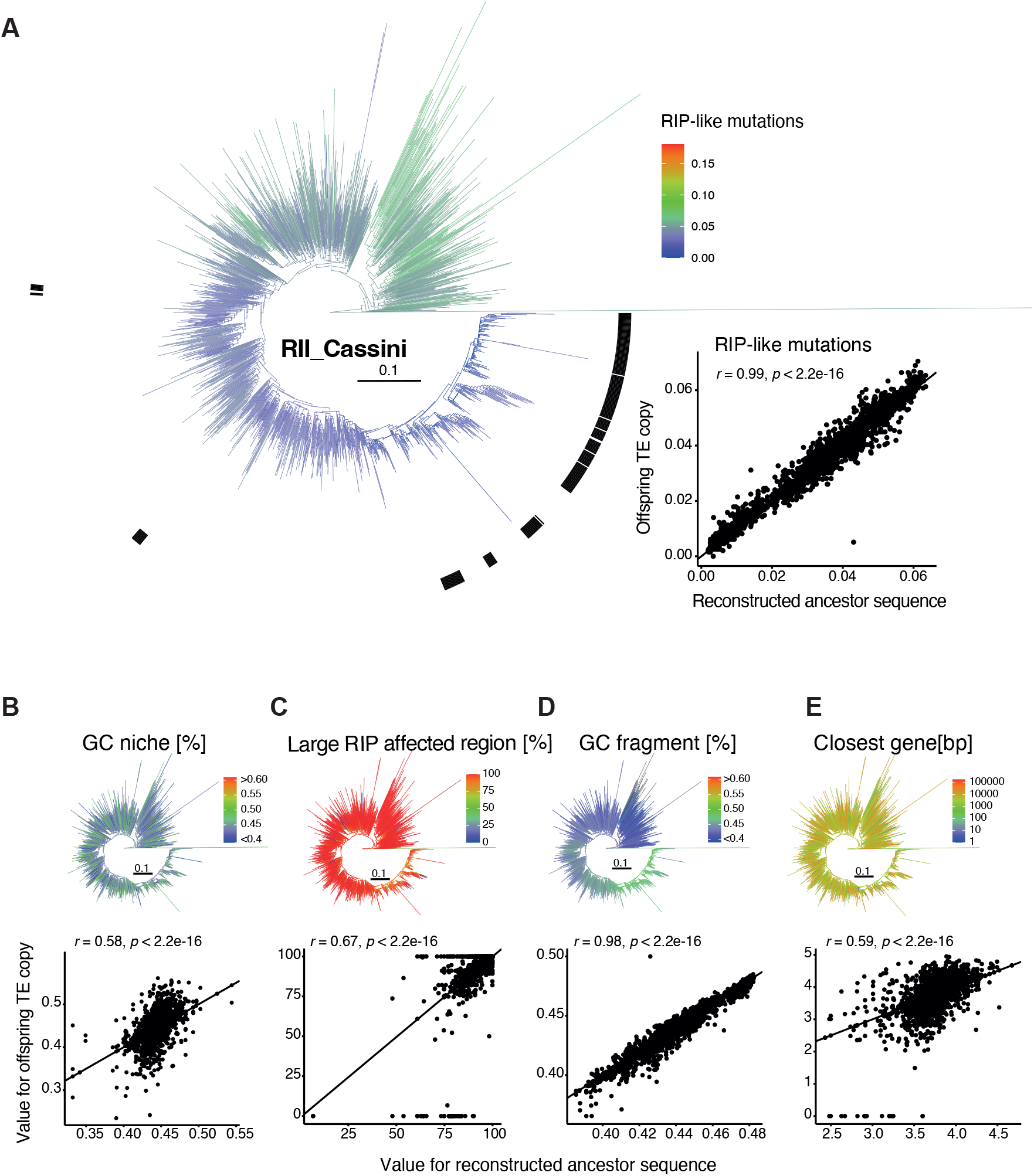
Phylogenetic tree of the *Cassini* retrotransposon: (A) Phylogenetic tree with colors indicating the number of RIP-like mutations. The black bar marks the different burst clades. The dot plot shows the changes in RIP-like mutations from the ancestor to offspring for all internal and terminal branches based on the ancestral state reconstruction. (B-E) Ancestor-offspring changes for (B) the GC content of the niche, (C) the overlap of the niche with large RIP affected regions (LRAR), (D) the GC content of the copy and (E) the distance to the closest gene.

**Figure 7:**
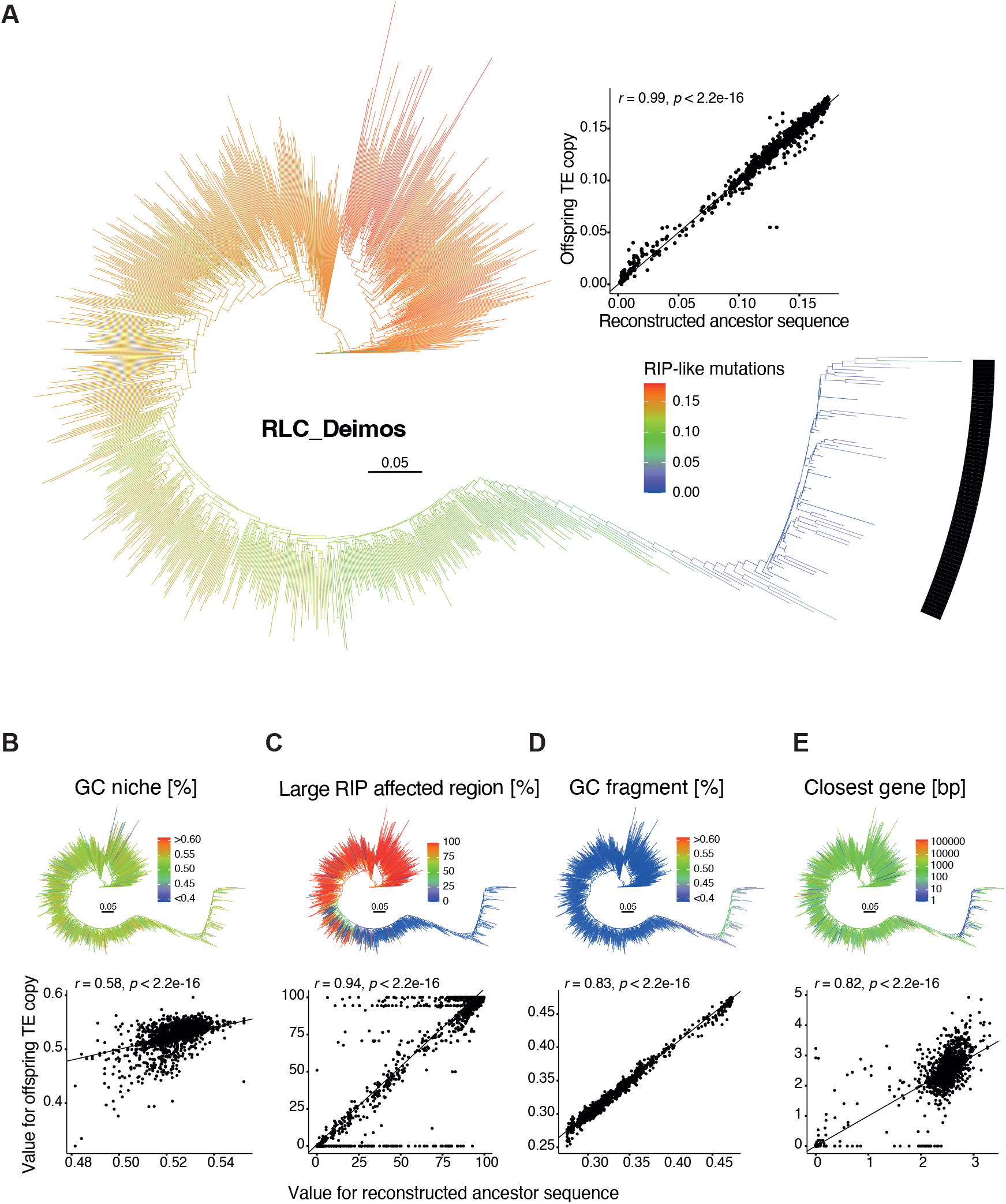
Phylogenetic tree of the *Copia* TE family *Deimos*: (A) Phylogenetic tree with colors indicating the number of RIP-like mutations. The black bar marks the different burst clades. The dot plot shows the changes in RIP-like mutations from the ancestor to offspring for all internal and terminal branches from the ancestral state reconstruction. (B-E) Phylogenetic trees and ancestor-offspring changes for (B) the GC content of the niche, (C) the overlap of the niche with large RIP affected regions, (D) the GC content of the copy and (E) the distance to the closest gene.

## DISCUSSION

TE activity is an important disruptor of genomic integrity and the potential for deleterious effects is strongly influenced by insertion site preferences. Our joint analyses of 19 complete *Z. tritici* genomes demonstrated that multiple TE families have been highly active in the recent evolutionary history of this species. A substantial number TEs have produced one or multiple bursts of proliferation that have distinct insertion site characteristics closer to genes and into regions with lower GC content compared with copies unrelated to the burst. Genomic defense mechanisms were only partially effective against preventing the proliferation of TEs. Our analyses indicate that a key trigger point for initiating new bursts is the successful insertion close to coding sequences. Hence, the genomic environment likely plays a key role in the evolutionary success of TE families.

We identified the emergence and expansion of numerous clades within TE families consistent with a burst of rapid copy number expansions. TE families often include large numbers of inactive copies that accumulate mutations and are unlikely to cause new insertions. Consistent with the dynamic nature of TE families, many TEs of *Z. tritici* are not detectable in the genomes of sister species. In particular, MITE families are most often absent in the sister species. This is consistent with rapid divergence of these non-autonomous elements devoid of coding sequences. Searching for more distant homologs to *Z. tritici* TEs might help identify additional putative ancestors, as observed in other species (Wicker *et al*., 2007; Bleykasten-Grosshans *et al*., 2021). Rapid proliferation of new TE copies is highly uneven among TE families, with a small subset of families driving the majority of recent insertions. Such new insertions have persisted despite potential purifying selection against new insertions, mutations introduced by genomics defenses (*i*.*e*., RIP) or silencing. TE families such as the RLC_Deimos were activated in several waves starting new branches or “subfamilies” that originated from distinct TE copies in the genome.

Active TEs that generate identical copies typically trigger defense mechanisms. Hence, such TE families are the most likely to be highly affected by RIP mutations. However, our results suggest RIP has only a weak impact on young TEs. Weak RIP signatures were predominantly found in MITEs. Short elements are most likely to escape detection by the RIP machinery as seen in another species (Pereira *et al*., 2021). Longer elements show stronger impacts of RIP using both GC content and RIP-like mutations as proxies. We found evidence for RIP mostly in older TE copies, while copies generated during recent bursts show nearly no RIP signatures. Escaping the effects of RIP may be a prerequisite for the initiation of a burst, hence the strong association of age and RIP mutations. RIP has mostly been studied in the Ascomycete *N. crassa*, where RIP introduces a large number of mutations after just one generation of sexual recombination (Wang *et al*., 2020b). The life cycle of *Z. tritici* is thought to consist of several cycles of asexual reproduction during the growing season, and only one round of sexual reproduction at the end of the season (Chen & McDonald, 1996). Hence, TEs may proliferate despite RIP for short periods but ubiquitous sexual reproduction should provide ample opportunity for RIP to act on copies of any recent burst. Evidence for RIP mutations are widespread in the genome of *Z. tritici* and the machinery for RIP is present in at least some isolates (van Wyk *et al*., 2021). Species-wide analyses have suggested that newly established populations of the pathogen have lost a functional RIP machinery (Lorrain *et al*., 2020). Loss of the RIP machinery would help preserve nearly identical copies of recently duplicated TE sequences and would maintain GC content at high levels. To what extent the loss of RIP has shaped the distribution of GC content across the genome including TEs copies remains unknown. The absence of RIP-like mutations in nearly all recently expanded TE families strongly suggests that recent TE proliferation was enabled by weakened genome defenses in the species.

A mechanistic understanding of triggers activating TEs is largely lacking, yet here we show the importance of the insertion niche. Young TE copies have distinct associations with particular genomic niches compared with older copies, which tend to be located in niches with high TE content. In contrast, copies triggering recent bursts and the resulting burst copies themselves tend to be inserted closer to genes. However, the association of young TE copies with genomic niches is most likely confounded by the action of purifying selection. As TEs can disrupt coding sequences or change expression profiles of neighboring genes, purifying selection is most likely strongest against TEs inserting into gene rich niches. In contrast, TEs inserting into TE-rich niches possibly causing nested insertions likely have only a minor impact on fitness. In some fungal plant pathogens including *Z. tritici*, such nested insertions led to the compartmentalization of niches with high TE density and niches mostly composed of genes (Plissonneau *et al*., 2016). Repeated insertions of TEs into such TE-rich niches likely exacerbated genome compartmentalization. Selection is also expected to act on the effectiveness of defenses against TEs. Silencing or hypermutation of TEs close to genes may disrupt the functionality of the genes as well; hence, the efficiency of genomic defenses against TEs may be weakened by selection. It is thus conceivable that otherwise silenced TEs can remain both functional and active when inserted close to a gene. Additionally, there are indications that RIP was lost in some isolates of *Z. tritici* (Möller *et al*., 2021), which would greatly facilitate TE activity across the genome.

Beyond the benefit of weaker genomic defenses, the propensity of bursts being initiated by TEs inserting near genes may also be related to beneficial impacts of the TE insertion itself. We found that copies at the start of bursts tend to have higher allele frequency within the species (*i*.*e*., present in most analyzed genomes). Larger population genomic datasets will enable the analysis of selection acting on parental copies initiating bursts. It is conceivable though that the beneficial effects of an individual TE insertion are linked to triggers of TE bursts. However, selection at the organismal level driven by fitness benefits conferred by specific TE insertions is compounded by TE-level selection for higher proliferation rates. Such multi-level selection was recently suggested to represent a Devil’s Bargain in plant pathogens trading short term benefits of TE insertions with longer term risks of genome expansions (Fouché *et al*., 2022). Assessing fitness effects of individual TE insertions along with their expansion history will enable further hypothesis testing about the proximate drivers of TE expansions over a range of evolutionary time scales.

## METHODS

### GENOME SEQUENCES AND TRANSPOSABLE ELEMENT DETECTION

We used a set of 19 reference-quality genomes of *Z. tritici* assembled using PacBio sequencing (Badet *et al*., 2020; European Nucleotide Archive BioProject PRJEB33986). The genomes cover the global genetic diversity of the species with isolates originating from 14 countries and six continents (Figure 1A, Supplementary Table S1). We used an improved TE annotation for the species with elements retrieved from all assembled genomes (Badet *et al*., 2020). TE annotation steps included using RepeatMasker, LTR-Finder, MITE-Tracker, SINE-Finder, Sine-Scan and extensive manual curation with WICKERsoft (Smit *et al*.; Xu & Wang, 2007; Breen *et al*., 2010; Wenke *et al*., 2011; Ma *et al*., 2015; Gao *et al*., 2016; Mao & Wang, 2017; Crescente *et al*., 2018; Badet *et al*., 2020). The primary TE annotation was followed by stringent filtering steps to detect nested insertions and to join TE fragments. Simple repeats, low complexity regions and elements smaller than 100 bp were removed. TEs belonging to the same family overlapping by more than 100 bp were merged. TEs belonging to different families overlapping by more than 100 bp were considered as nested insertions. TEs belonging to the same family separated by less than 200 bp were considered as fragmented TEs and merged into a single element (Badet *et al*., 2020). We additionally annotated TEs using the same pipeline in high quality genomes of the sister species *Z. ardabiliae, Z. brevis, Z. pseudotritici* and *Z. passerinii* (Feurtey *et al*., 2020).

### MULTIPLE SEQUENCE ALIGNMENTS

We created multiple sequence alignments for all copies belonging to the same TE family from the 19 *Z. tritici* and four sister species genomes (Supplementary Figure S4). We extracted all sequences of TE families with copy numbers ≥ 20 with the function *faidx* in samtools version 1.9 (Li *et al*., 2009). In case of fragmented elements, we extracted all fragments as individual copies. We reverse-complemented sequences where necessary prior to sequence alignment. To extract coding regions, we performed blastx searches against the PTREP18 database and against the non-redundant protein database from NCBI (09/2020) with diamond blast version 0.9.32.133 and selected the hit with the highest bit score with at least 200 bp length (Thomas Wicker; http://botserv2.uzh.ch/kelldata/trep-db/index.html) (Altschul *et al*., 1990; Buchfink *et al*., 2014). For small non-autonomous TE families lacking a coding region, we retained the entire sequence. We created multiple sequence alignments for each family with MAFFT version 7.453 and the following parameters: --thread 1 --reorder --localpair - -maxiterate 1000 --nomemsave --leavegappyregion (Katoh & Standley, 2013). For four TE families with high copy numbers and large coding regions (RII_Cassini, RLG_Luna, RLG_Sol, RLC_Deimos), we slightly decreased accuracy of MAFFT, using the parameters --6merpair instead of --localpair.

### TE FAMILY DIVERGENCE

TE families are expected to be active during different time spans and evolve at different rates. To estimate the relative age of the TE families, we ran RIPCAL with --windowsize 1000 --model consensus to create an additional consensus sequence that includes all copies of a TE family (Hane & Oliver, 2008, 2010). In R version 4.0.2 we created DNAbin objects with the R package ape version 5.3 and calculated nucleotide diversity of the multiple sequence alignments for each TE family with *nuc*.*div* in the package pegas version 0.13 (Paradis, 2010; Paradis & Schliep, 2019; R Core Team, 2020). To compare between TE families, we divided the nucleotide diversity by the length of the corresponding TE coding region. We estimated the relative age of TE bursts per family using RepeatMasker. To compare recent activity or bursts, we created a repeat landscape using *build Summary, calcDivergenceFromAlign* using Kimura divergence and *createRepeatLandscape* in RepeatMasker and visualized the results with ggplot (Kimura, 1980).

### GENOMIC ENVIRONMENT OF TE INSERTIONS

We described the genomic characteristics of niches containing TE insertions. For 5 kb niches centered around TE insertion loci, we calculated the TE and gene content based on TE and gene annotations, respectively, using the *intersect* command in bedtools version 2.28.0. We calculated GC content with the *geecee* tool in EMBOSS version 6.6.0 (Rice *et al*., 2000; Quinlan & Hall, 2010; Grandaubert *et al*., 2015). We also calculated the distance to the closest gene and TE with the *closest* command in bedtools. We used Occultercut version 1.1 with default parameters to detect isochores with low (≤ 49%) or moderate (> 49%) GC content (Testa *et al*., 2016). We used TheRIPper to identify large RIP affected regions in all analyzed genomes, and calculated the overlap of TE insertions and RIP affected regions with bedtools *intersect* (van Wyk *et al*., 2019). For the reference genome IPO323, we used available ChIP-seq information (http://ascobase.cgrb.oregonstate.edu/cgi-bin/gb2/gbrowse/ztitici_public/) to define the chromatin structure in niches around TE insertions (Schotanus *et al*., 2015).

### CHARACTERISTICS OF TE INSERTIONS

Many TE sequences in the genome are fragmented due to nested insertions or partial deletions. To improve the quality of multiple sequence alignments, we selected only TE coding regions for phylogenetic analyses. For this, we trimmed the multiple sequence alignments with *extractalign* from EMBOSS, based on the position of the coding sequences, removed empty sequences with trimAl -gt 0 version 1.4.rev15 (http://trimal.cgenomics.org) and removed fragments that contained more than 50% gap positions in the coding region (Capella-Gutiérrez *et al*., 2009). We calculated the GC content of each TE coding region with *geecee* in EMBOSS. To quantify RIP-like mutation signatures, we extracted dinucleotide frequencies for each TE family alignment with *count* in the package seqinr (Charif & Lobry, 2007). To define locus specific TE dynamics, we identified first the closest up- and downstream fixed orthologous genes based on the annotation of the pangenome with *closest* in bedtools (Badet *et al*., 2020). Next, we defined TE insertions belonging to the same TE family and being located between the same fixed orthologous genes as orthologous TE groups. Visualizations were made with ggplot (Wickham, 2016).

### MAXIMUM LIKELIHOOD TREES

We estimated maximum likelihood trees for all TE families with indications for recent activity and bursts in the species. We extracted conserved blocks of the coding region with Gblocks version 0.91b, using the following parameters: -t=d -b3=10 -b4=5 -b5=a -b0=5 (Castresana, 2000). For each TE family, we included two sequences retrieved from the same TE in sister species genomes to root trees. We estimated maximum likelihood trees with RAxML version 8.2 (Stamatakis, 2014). For this, we generated 20 ML trees with each a different starting tree and extracted the starting tree with best likelihood with the following parameters: raxmlHPC-PTHREADS-SSE3 -T 4 -m GTRGAMMA -p 12345 -# 10 --print-identical-sequences. We performed a bootstrap analysis to obtain branch support values with the following parameters: raxmlHPC-PTHREADS-SSE3 -T 4 -m GTRGAMMA -p 12345 -b 12345 -# 50 --print-identical-sequences. Finally, we added bipartitions on the best ML tree with the following parameters: raxmlHPC-PTHREADS-SSE3 -T 4 -m GTRGAMMA -p 12345 -f b --print-identical-sequences.

### ANCESTRAL STATE RECONSTRUCTION

We performed ancestral state reconstruction for each TE family including characteristics of the TE sequences or the niche of the TE insertion. We imported the trees into R using *read*.*tree* from the package treeio version 1.10.0 (Wang *et al*., 2020a). We rooted trees with *root* in the R package ape, using sister species sequences as outgroups. We converted tree objects to tibble with *as_tibble* from the package tibble version 3.0.1 in tidyverse version 1.3.0 and added metadata using dplyr version 0.8.5 in tidyverse (Wickham *et al*., 2019, 2020; Müller & Wickham, 2020). We converted the tibble objects back to tree formats with *as*.*treedata* in treeio (Wickham *et al*., 2019). Using *fastAnc* and *contMap* from package phytools version 0.7-47, we performed ancestral state reconstruction for characteristics of the following continuous traits: gene density, TE density and GC content of 2.5 kb niches surrounding the TEs, closest gene, GC and RIP-like mutations per bp (Revell, 2012). To estimate ancestral states for binary characteristics (GC rich *vs*. poor, core *vs*. accessory chromosome), we used *make*.*simmap* from the package phytools with an equal rates model and 100 simulations. We visualized the trees with *ggtree* version 2.0.1 (Yu *et al*., 2017). To retrieve clades representing recent bursts, we created polytomy trees from binary trees, using the command *CollapseNode* from TreeTools version 1.4.4 in R at branch lengths smaller than 1.1e-05 (Smith, 2019). For each burst clade, we defined the parental branch and an outgroup of the clade with *offspring* in the *treeio* package as the ancestral branch. We compared niche metrics of ancestral branches of bursts with the distribution of the same metric in all elements outside of bursts. We performed associating mapping for shared characteristics along the phylogenetic tree using treeWAS (Collins & Didelot, 2018).

## ACKNOWLEDGEMENTS

We are very grateful for helpful comments on an earlier version of the manuscript by Emile Gluck-Thaler, for discussions about phylogenetic inference with Vinciane Mossion and statistical advice by Claudia Sarai Reyes-Avila. DC is supported by the Swiss National Science and the Fondation Pierre Mercier pour la Science.

