## Supplementary Figure for "Recent transposable element bursts triggered by insertions near genes in a fungal pathogen"

### Supplementary figures

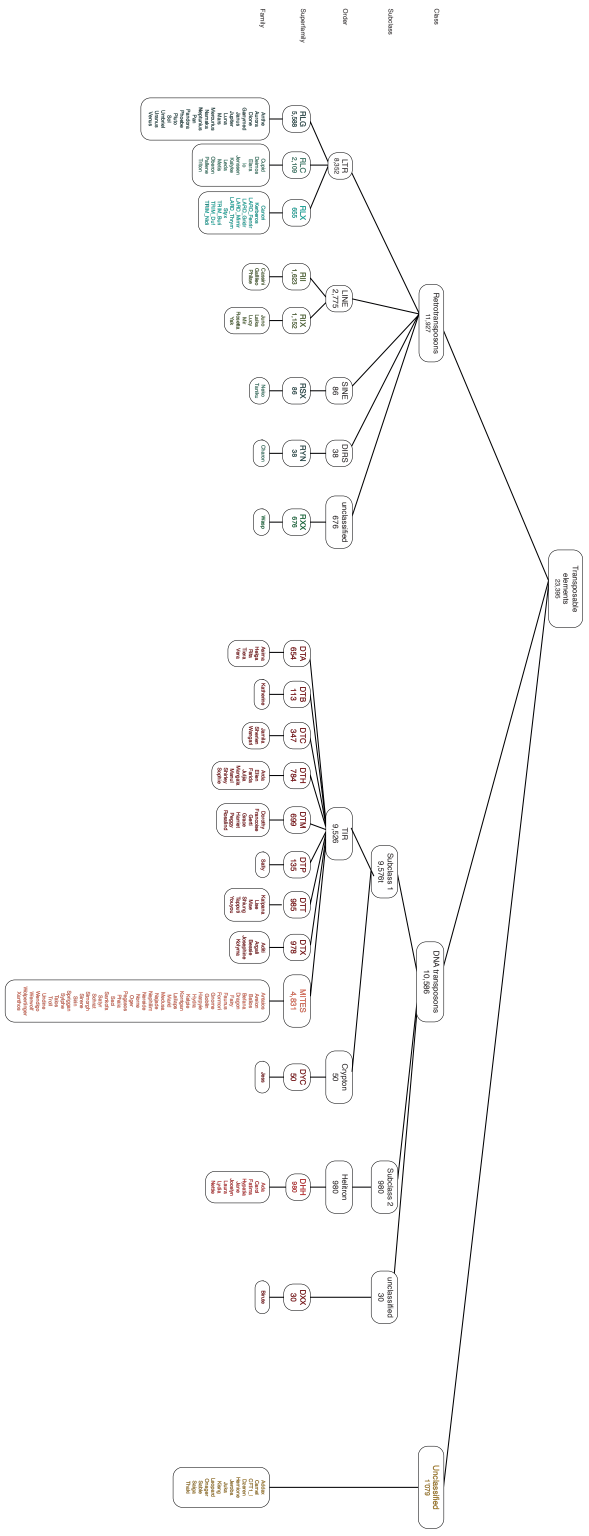

**Supplementary Figure S1: Hierarchy TE superfamilies**: Classes, subclasses, orders, superfamilies as well as the tree-letter code according to Wicker et al (2007). The *Z. tritici* specific family names are according to Badet et al (2020).

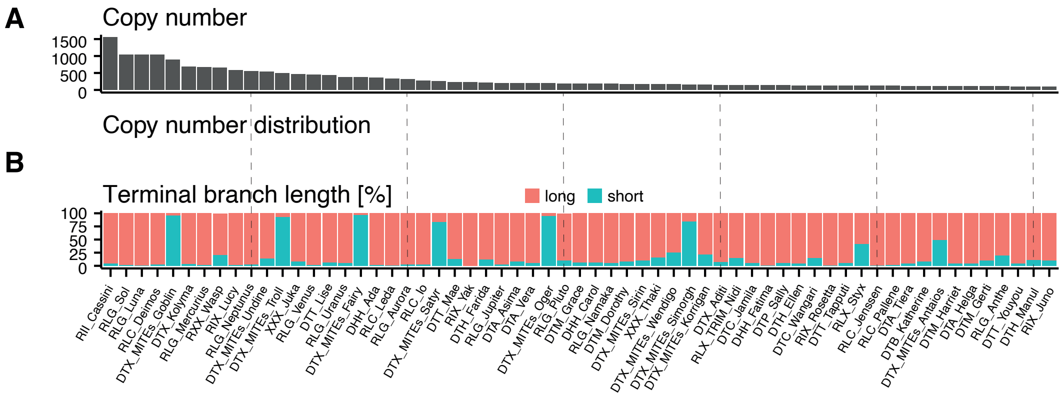


**Supplementary Figure S2: Characteristics of high-copy TE families**: TE families are ordered from the highest copy numbers to lowest copy numbers (right) in all 19 analyzed genomes combined. (A) Total copy numbers. (B) Long (> 0.00001; red) and short (≤0.00001; blue) terminal branch lengths of individual copies characterizing two classes of divergence times.

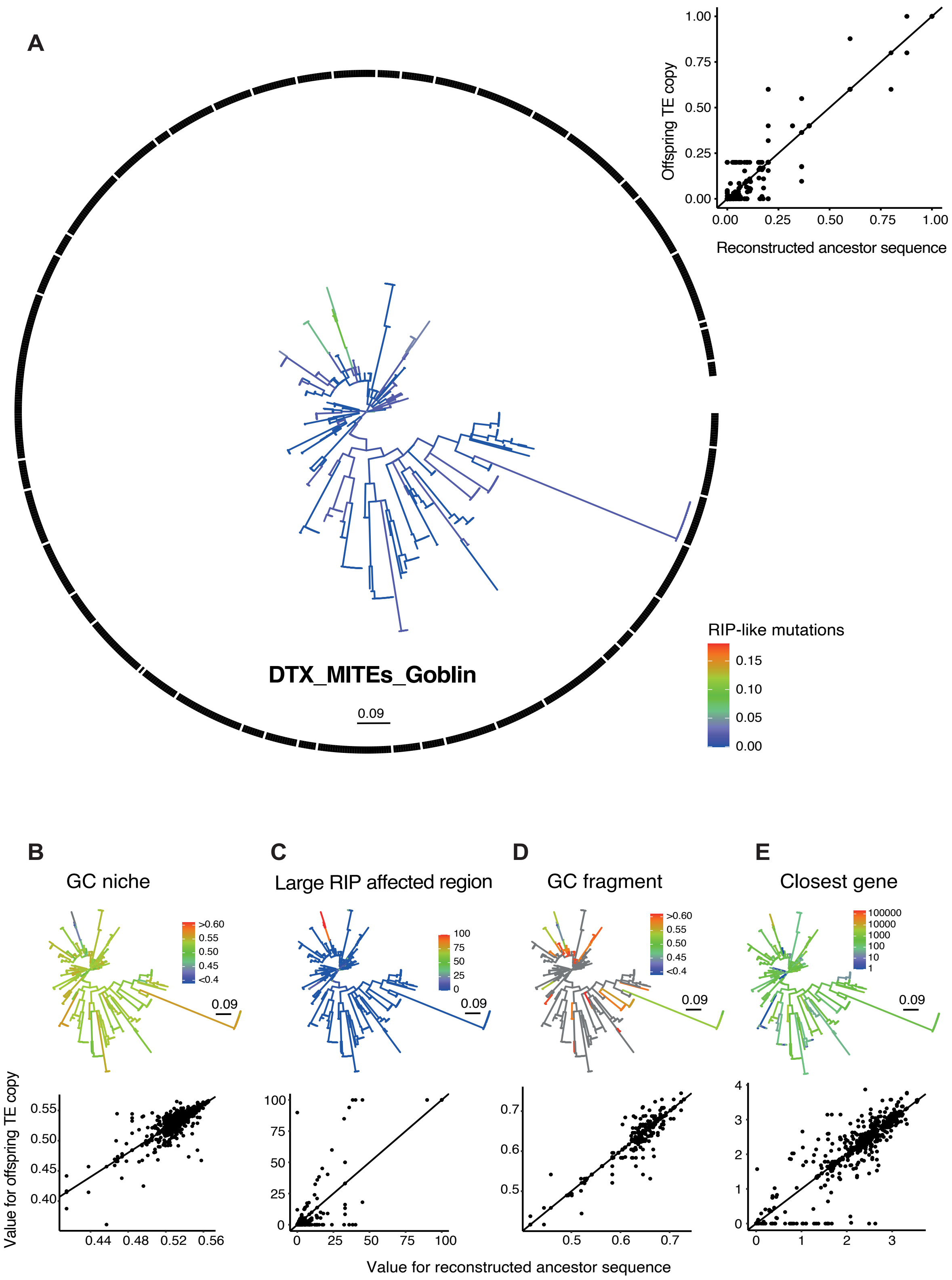


**Supplementary Figure S3: Phylogenetic tree of the TE family DTX_MITE_Gobblin**: (A) Phylogenetic tree with colors indicating the number of RIP-like mutations. The black bar marks the different burst clades. The dot plot shows the changes in RIP-like mutations from the ancestor to offspring for all internal and terminal branches from the ancestral state reconstruction. (B-E)

Phylogenetic trees and ancestor-offspring changes for (B) the GC content of the niche, (C) the overlap of the niche with large RIP affected regions, (D) the GC content of the copy and (E) the distance to the closest gene.

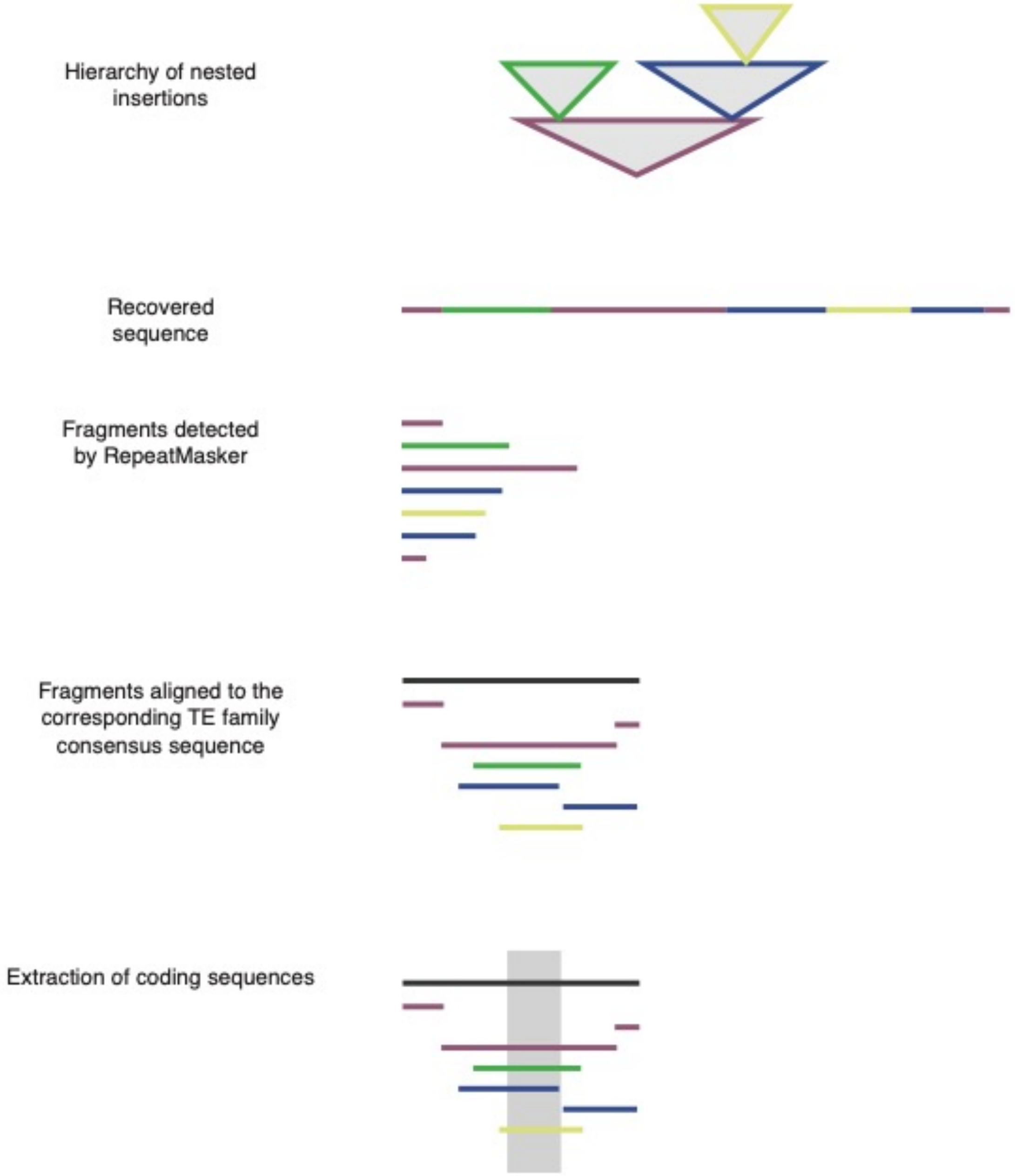


**Supplementary Figure S4: Procedure to obtain multiple sequence alignments among copies of TE families**. Due to the high number of nested insertions and partially deleted fragments, we aligned only coding regions.

#### Supplementary tables

(see separate xls file)

**Supplementary Table S1**: Collection of 19 reference-quality genomes of *Zymoseptoria tritici*. Data from Badet et al (2020)

**Supplementary Table S2**: Metadata for all TE insertion loci. Includes TE family, information about isolates of origin, position in the genome, niche and TE sequence characteristics.
